# Pervasive patterns in the songs of passerine birds resemble human music universals and are linked with production and cognitive mechanisms

**DOI:** 10.1101/2024.07.15.603339

**Authors:** Logan S James, Kendra Oudyk, Erin M Wall, Yining Chen, William D Pearse, Jon T Sakata

**Affiliations:** Department of Biology, McGill University, Montréal, Quebec, Canada; Department of Integrative Biology, University of Texas at Austin, Austin, TX, USA; Integrated Program in Neuroscience, McGill University, Montréal, Quebec, Canada; Department of Life Sciences, Imperial College London, Silwood Park Campus, Ascot, Berkshire, UK; Centre for Research in Brain, Language and Music, McGill University, Montréal, Quebec, Canada

## Abstract

Music is a complex learned behavior that is ubiquitous among humans, and many musical patterns are shared across geography and cultures (“music universals”). Knowing whether these universals are specific to humans or shared with other animals is important to understand how production-related factors (motor biases and constraints) or cognitive factors (learning) contribute to the emergence of these acoustic patterns. Bird song is often described as an animal analogue of human music, and some studies of individual avian species highlight acoustic similarities between bird song and music. However, expansive and comparative approaches are necessary to identify universal patterns within bird song, reveal mechanisms associated with these patterns, and draw parallels to music universals. Here, we adopt such an approach and analyze the prevalence of acoustic patterns (sequences) across ∼300 species of passerines, spanning both oscines (songbirds; vocal learners) and their sister clade, suboscines (passerines that produce songs that are not learned), as well as within a global corpus of human vocal music. This approach allowed us to directly test hypotheses that phonation mechanisms or vocal learning shape the emergence of universal patterns. We first document acoustic patterns that were widely shared across passerines and similar to music universals (e.g., small pitch intervals), highlighting the role of shared vocal production mechanisms in these patterns. Consistent with a contribution of vocal learning, we observed patterns (e.g., alternation in durations) there were more similar between oscines and humans than between suboscines and humans. Interestingly, we also discovered patterns (e.g., pitch alternation) that were inconsistent with a contribution of vocal learning and were more similar between suboscines and humans than between oscines and humans. This research provides the broadest evidence of shared universals in vocal performance across birds and humans and highlights convergent mechanisms shaping communication patterns.

## Introduction

The complex vocalizations of birds are referred to as “songs” across many human languages (Marler & Slabbekoorn, 2004; Baptista & Keister, 2005). Indeed, there are numerous parallels between bird song and human music, with both consisting of sequences of acoustic elements that are temporally ordered and hierarchically organized (Marler & Slabbekoorn, 2004; Patel, 2010; Rothenberg *et al*., 2014; Fitch, 2015; Honing *et al*., 2015; Rohrmeier *et al*., 2015). Both bird song and music rely on complex motor coordination for production and auditory processing mechanisms that induce physiological changes in receivers (Marler & Slabbekoorn, 2004; Woolley & Doupe, 2008; Byers, Hebets & Podos, 2010; Elemans *et al*., 2015; Schmidt & Goller, 2016; Sakata & Yazaki-Sugiyama, 2020; Bainbridge *et al*., 2021). Further, music and bird song vary according to social context and target different receivers (Woolley & Doupe, 2008; Keller, Novembre & Hove, 2014; Chen, Matheson & Sakata, 2016; Mehr *et al*., 2018; Coleman *et al*., 2021).

Structurally, music around the world shares “universal” features, defined as patterns that are more prevalent than expected by chance (Savage *et al*., 2015; Mehr *et al*., 2019). For example, differences in pitch between consecutive notes in a song are smaller than expected by chance (small pitch intervals), and notes at the end of musical phrases tend to be longer in duration than expected by chance (phrase-final elongation; (Tierney, Russo & Patel, 2011; Savage *et al*., 2015; Mehr *et al*., 2019). Furthermore, compared to all possible pitch contours, descending and arched contours appear more often than expected (Huron, 1996; Tierney *et al*., 2011; Savage *et al*., 2015). However, the mechanistic and evolutionary origins of these patterns remain largely unknown.

Uncovering the degree to which these music universals are shared with non-human animals will lend crucial insight into the biological origins of these patterns and will generate hypotheses about convergent neural and anatomical mechanisms (Fitch, 2006; Patel, 2010; Rothenberg *et al*., 2014; Hoeschele *et al*., 2015; Podos & Sung, 2020). While previous studies identify some vocal patterns that are similar between humans and non-human animals (Dobson & Lemon, 1977; Tierney, Russo & Patel, 2008; Tierney *et al*., 2011; Araya-Salas, 2012; Doolittle *et al*., 2014; Bregman, Patel & Gentner, 2016; Roeske, Kelty-Stephen & Wallot, 2018), no study has investigated these patterns across the breadth and diversity of species within a taxonomic group required to make broad inferences about similarity.

Here, we use a phylogenetic approach to examine the prevalence of music universals within the songs of ∼300 passerine species. The avian order Passeriformes (passerines) is a taxonomically rich order that consists of oscines (vocal learners) and suboscines (vocal non-learners), offering a powerful opportunity for an expansive investigation into potential mechanisms underlying music universals (e.g., vocal learning and/or biases or constraints in vocal production). We analyze identical patterns in a corpus of global human song to directly compare the magnitudes of these universals across birds and humans.

Given that bird song and human music both rely on the coordination of vocal musculature and breathing and share various production mechanisms (Riede & Goller, 2010; Tierney *et al*., 2011; Elemans *et al*., 2015; Schmidt & Goller, 2016; Podos & Sung, 2020; Goller, 2022a), we anticipate a number of common patterns to be consistent across human and passerine songs because of broadly shared production mechanisms. However, because passerines and humans differ in their breathing patterns during singing and use different structures for sound production (Schmidt & Goller, 2016; Goller, 2022a), we expect some patterns that are widely prevalent within passerines to be distinct from those in humans. Furthermore, because music and oscine song are learned and culturally transmitted (in contrast to suboscine song), acoustic patterns in music that are more similar to oscine song than to suboscine song could result from shared cognitive or sensorimotor processes subserving vocal learning.

## Results

We curated, annotated, and analyzed a library of songs consisting of >64,000 syllables (continuous epochs of sound) from 299 species within 152 genera and 50 families of passerines (Table S1). In the first set of analyses, we reveal acoustic patterns that are widespread across the songs of passerines, targeting acoustic features commonly measured in music and patterns that are prevalent in music (frequency jumps, positional variation, and alternation; Figure 1). In the second set of analyses, we disentangle phylogenetic variation in these patterns, focusing on variation between oscines and suboscines. Finally, we analyze these same patterns in a dataset of world music and reveal similarities and differences across oscine, suboscine, and human songs.

**Figure 1.**
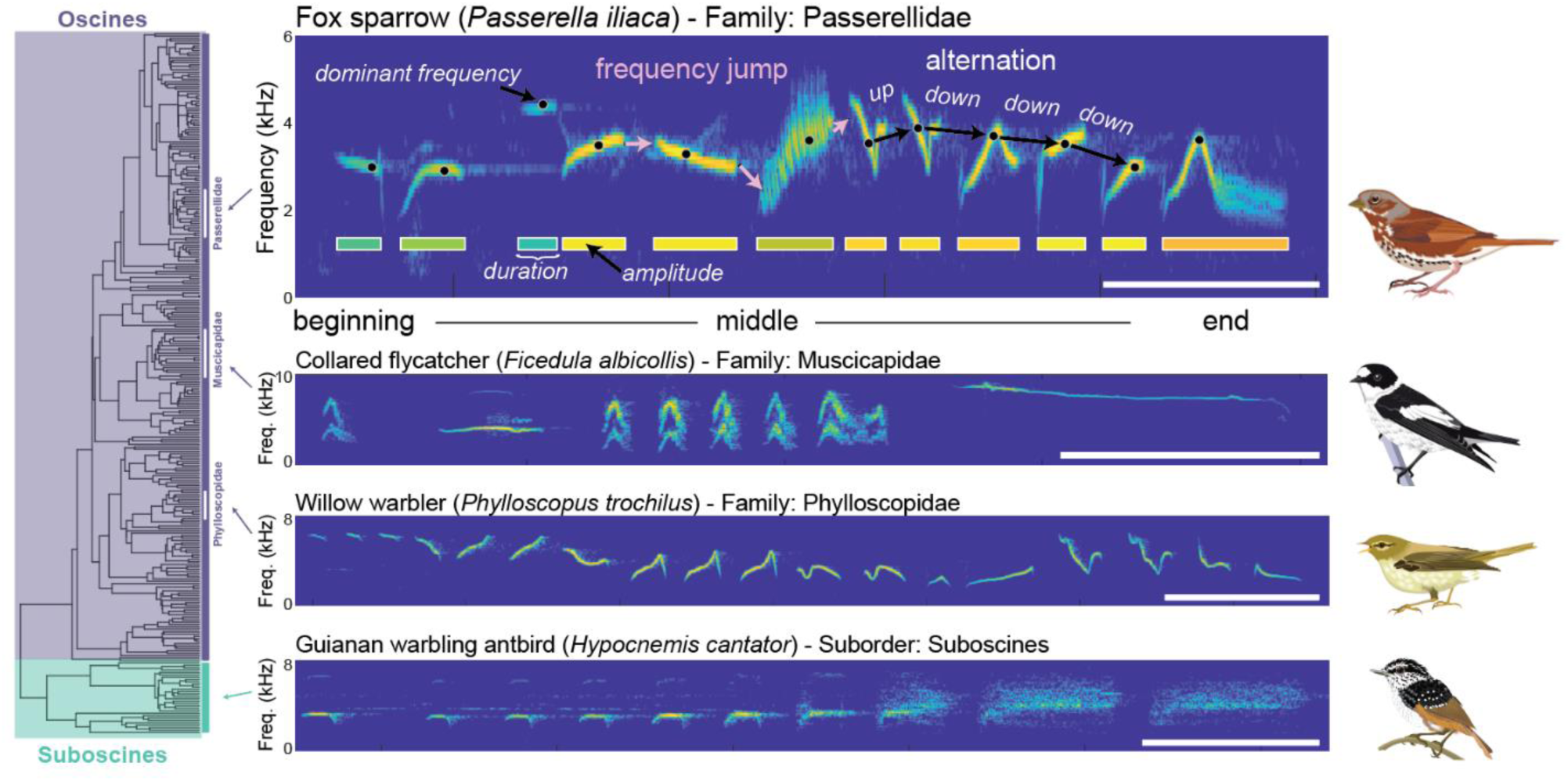
Analysis of acoustic features in the songs of passerines. Plotted are spectrograms (frequency on the y-axis, time on the x-axis, color as amplitude). Top: a spectrogram of one rendition of song of a fox sparrow (Passerella iliaca) with annotations depicting the acoustic features we measured and analyses we conducted. Pink arrows depict our analysis of frequency jumps comparing the dominant frequency at the end (final 8 ms) of a syllable to the frequency at the beginning (initial 8 ms) of the next syllable. Black dots on each syllable indicate the dominant frequency of that syllable, and black arrows illustrate our analysis of alternation of frequency. Rectangles below each syllable indicate the duration (length) and amplitude (color) of each syllable. For positional variation, we compared feature values from across the beginning, middle, and end syllables. Note that all annotations are approximate and used for illustrative purposes only. Below: example spectrograms from two additional oscine species: collared flycatcher (Ficedula albicollis) and willow warbler (Phylloscopus trochilus) as well as one suboscine species: Guianan warbling antbird (Hypocnemis cantator). Recordings provided (from top to bottom) by Patrik Åberg, Mikael Litsgård, Tony Fulford, and Jon King. Accessible at xeno-canto.org/406249, xeno-canto.org/288103, xeno-canto.org/331211, and xeno-canto.org/114053. Illustrations by Damond Kyllo. Scale bar= 0.5 sec.

### Frequency jumps in passerines

In music, pitch changes across adjacent notes are generally smaller than expected by chance (“small pitch intervals”; Huron, 1996; Savage *et al*., 2015; Mehr *et al*., 2019; Brinkman & Huron, 2021). We computed the average frequency jump across consecutive syllables (absolute value of the frequency difference from the end of one syllable to the beginning of the next syllable; Figure 1) for each passerine species with at least 5 consecutive syllables (*n* = 293 species; see Methods; mean ± SD: 16.1 ± 10.0 songs per species) and then compared this observed frequency jump to the expected frequency jump for that species (as computed through permutations of sequence randomization; see Methods). Frequency jumps in passerine songs were significantly smaller than expected by chance (MCMCglmm: Bayesian p < 0.001; Figure 2; Table S2).

**Figure 2:**
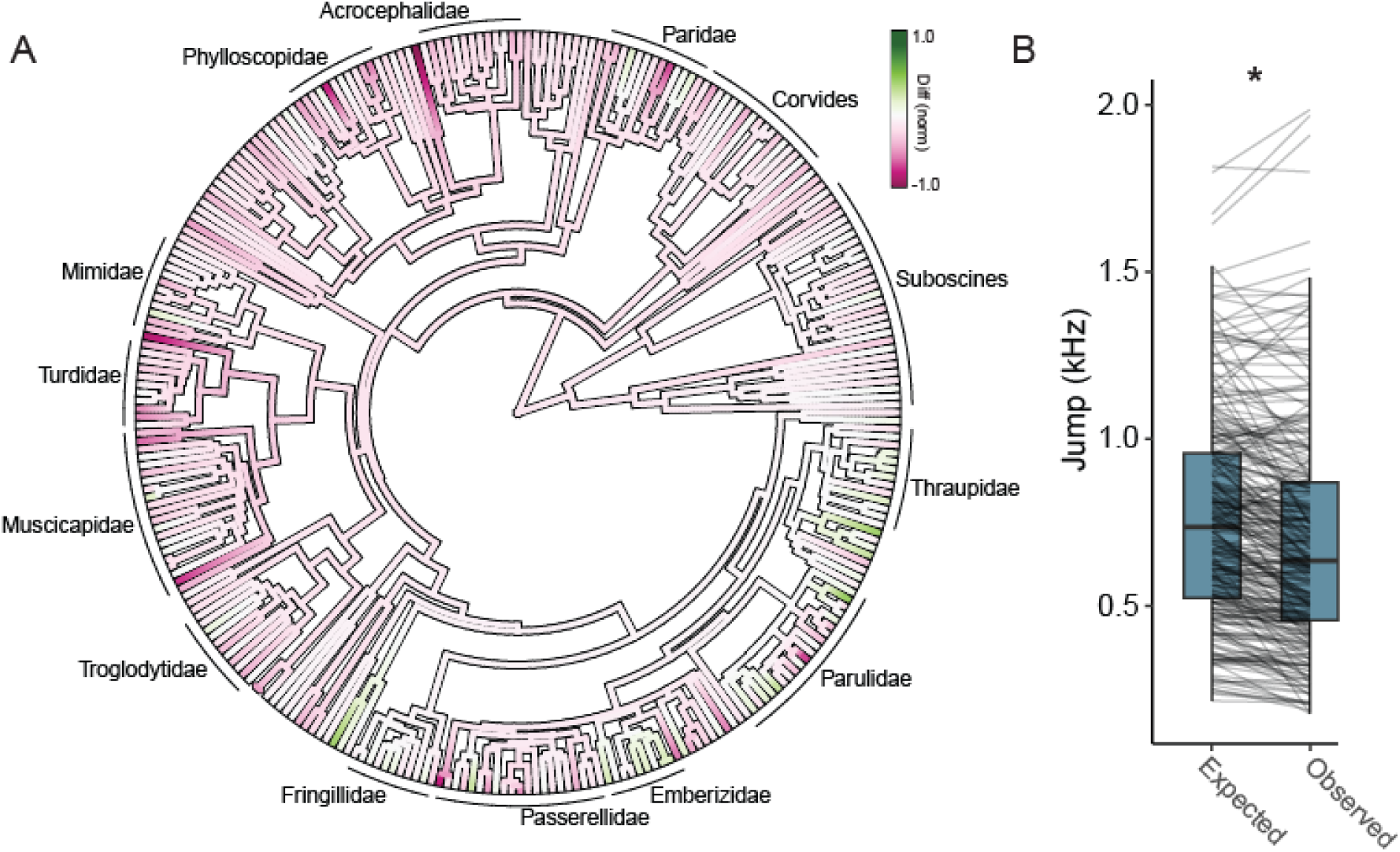
Frequency jumps in passerine song. A. Phylogenetic tree depicting the observed mean frequency jump minus the expected frequency jump (difference from expected) for each species. Pink colors indicate species where the change in frequency from the end of one syllable to the beginning of the next syllable tended to be smaller than expected (similar to music), whereas green colors indicate changes that were larger than expected. B. Box plot showing the overall distribution of the expected and observed frequency jumps, with lines connecting the data for a single species (each line depicts the values for an individual species).

### Positional variation in passerines

Common patterns of variation in the pitch, duration, and amplitude of acoustic units are observed in musical sequences (e.g., arched or descending pitch contours, phrase-final lengthening; (Palmer, 1989; Huron, 1996, 2006; Tierney *et al*., 2011; Savage *et al*., 2015; Kragness & Trainor, 2016; Mehr *et al*., 2019). We analyzed variation in acoustic features across syllables occupying beginning, middle, or end positions in the song (see Methods), and found significant positional variation in dominant frequency, duration, and amplitude that resembled patterns found in music (*n* = 292 species with data for all three positions; phylogenetically corrected MCMCglmm analyses; Figure 3; Table S2; mean ± SD: 16.2 ± 10.0 songs per species). Broadly speaking, (a) positional variation in dominant frequency resembled descending pitch contours (Figure 3A,B), (b) positional variation in syllable durations resembled phrase-final elongation (Figure 3C,D), and (c) positional variation in amplitude followed an arch-like contour (Figure 3E,F; Table S2).

**Figure 3.**
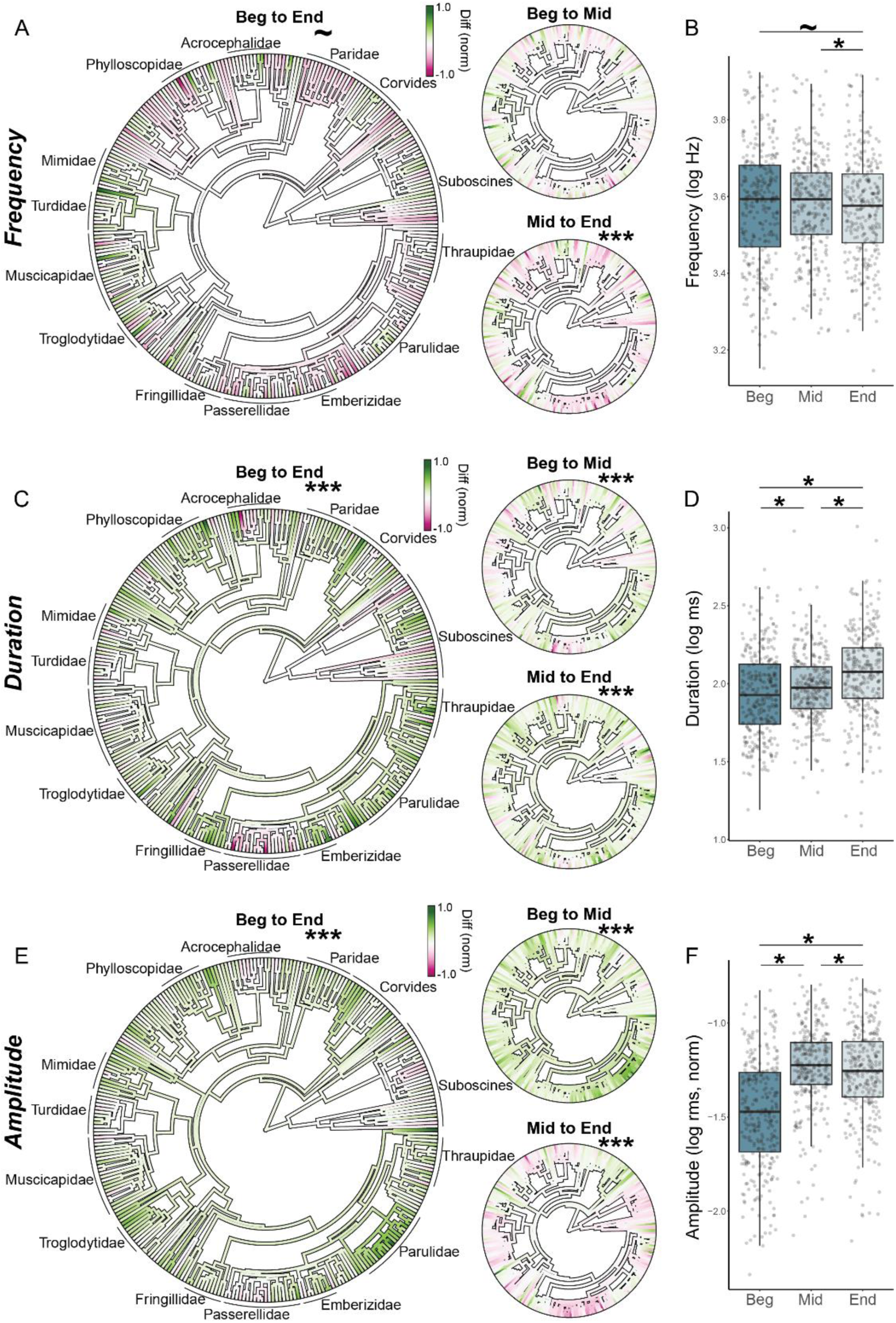
Positional variation in dominant frequency, duration, and amplitude. (A) Phylogenetic trees depicting the normalized difference in syllable dominant frequency across positions. Green colors indicate an increase across positions within the song, while pink colors indicate a decrease across positions. White indicates that the syllables were of the same values across positions. The large phylogeny on the left depicts the change from the beginning to the end yllable, the smaller phylogenies to the right depict the change from the beginning to the average middle syllable (top) and from the average middle to the end syllable (bottom). (B) Box plot showing the overall distribution of the data for each species (each dot depicts the average value for an individual species). Asterisks indicate significant differences based on phylogenetically corrected MCMCglmm analyses (Table S2). (C-F) Data plotted the same as (A-B) but for duration (C-D) and amplitude (E-F).

### Alternation in passerines

Music contains melodic structure in which, for example, the pitch of notes can rise and then fall across adjacent notes (alternation) or sequentially increase or sequentially decrease (no alternation; (Lerdahl & Jackendoff, 1983; Drake & Palmer, 1993; Patel, 2008). We characterized alternation in acoustic features across adjacent syllables and compared the observed prevalence of alternation to that observed in randomized sequences (chance). The prevalence of alternation in dominant frequency, duration, and amplitude were all significantly lower than expected by chance (*n* = 293 species; MCMCglmm: *p* < 0.001 for all; Figure 4; Table S2; mean ± SD: 16.1 ± 10.0 songs per species).

**Figure 4:**
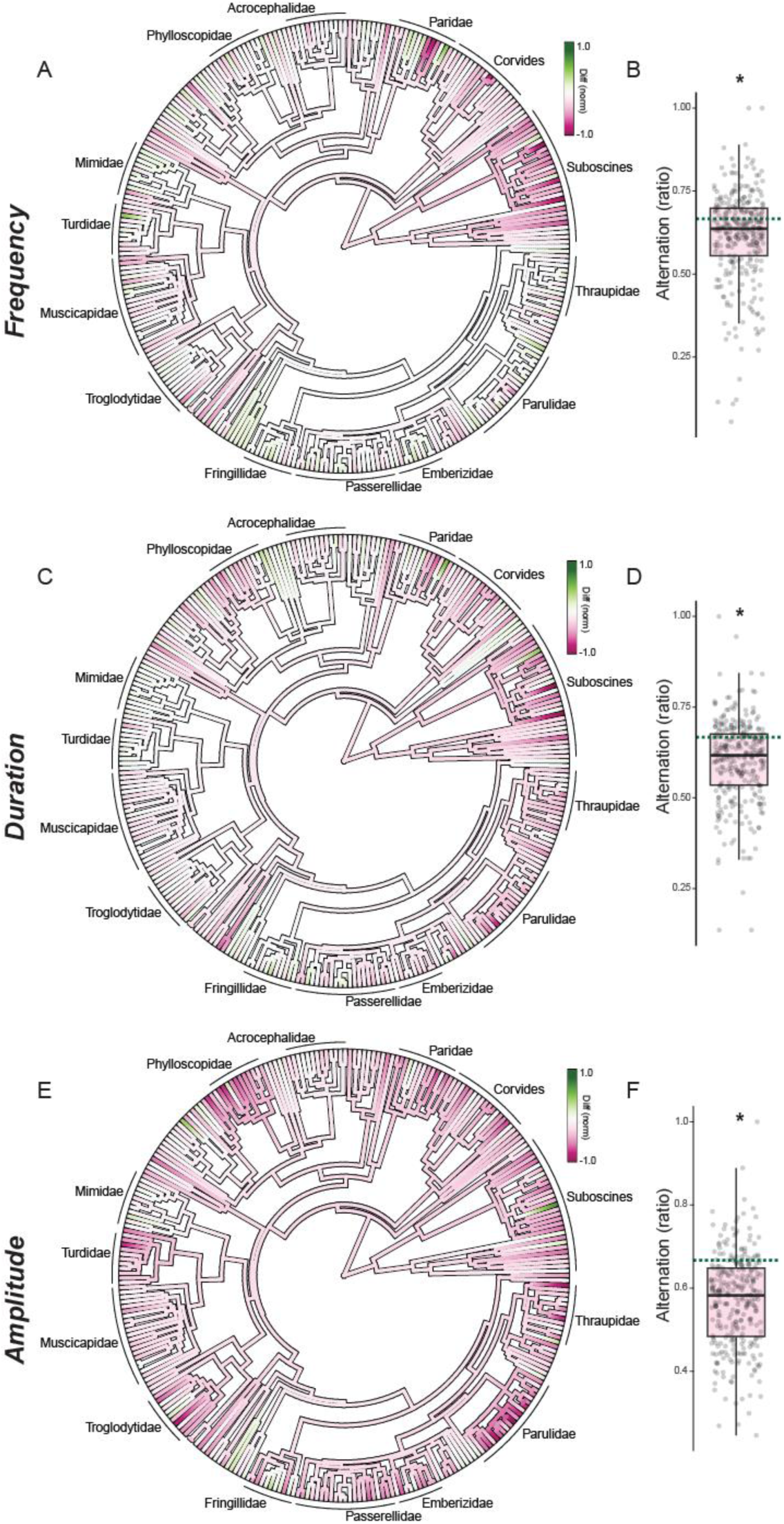
Alternation in passerine song. Phylogenetic trees (A, C, E) depict the difference of the observed prevalence of alternation (proportion of sequences with alternation) from chance (see Methods). Pink colors indicate species where alternation occurred less than expected by chance (similar to music), whereas green colors indicate more alternation than expected. Boxplots (B, D, F) depict the overall distribution of the observed prevalence of alternation for each species (each dot depicts the data from a single species), and dashed lines indicate chance.

### Comparison of acoustic patterns between oscines and suboscines

Passerines consist of oscines and suboscines, and these clades (suborders) demonstrate similarities as well as differences in many attributes, including morphology, neurobiology, and behavior (Gahr, 2000; Seddon, 2005; Amador, Goller & Mindlin, 2008; Derryberry *et al*., 2012; Liu *et al*., 2013; Schmidt & Goller, 2016; Garcia *et al*., 2017; Goller, Love & Mindlin, 2021; Amador & Mindlin, 2023). One major difference between these suborders is that oscines (songbirds) generally learn their vocalizations whereas suboscines produce vocalizations that are not learned (Kroodsma, 1984; Kroodsma & Konishi, 1991; Touchton, Seddon & Tobias, 2014; Jarvis, 2019; Ten Cate, 2021). As such, comparisons between oscines and suboscines could provide insight into the contribution of vocal learning (and its associated neural and morphological adaptations) to acoustic patterning. Furthermore, shared patterns across oscines and suboscines could highlight the role of shared motor production mechanisms.

Broadly speaking, frequency jumps were smaller than expected for both oscines and suboscines (Figure 5A). When analyzing both observed and expected jumps across clades, we continued to find that observed frequency jumps were significantly smaller than expected (*p* < 0.001), and furthermore that the degree of deviation from expected was not different between oscines and suboscines [interaction between clade (oscine or suboscine) and the observed versus expected frequency jump: *p* = 0.333; Figure 5A; Table S3]. We also observed a marginally significant main effect of clade (*p* = 0.069), with suboscines generally having smaller frequency jumps.

**Figure 5:**
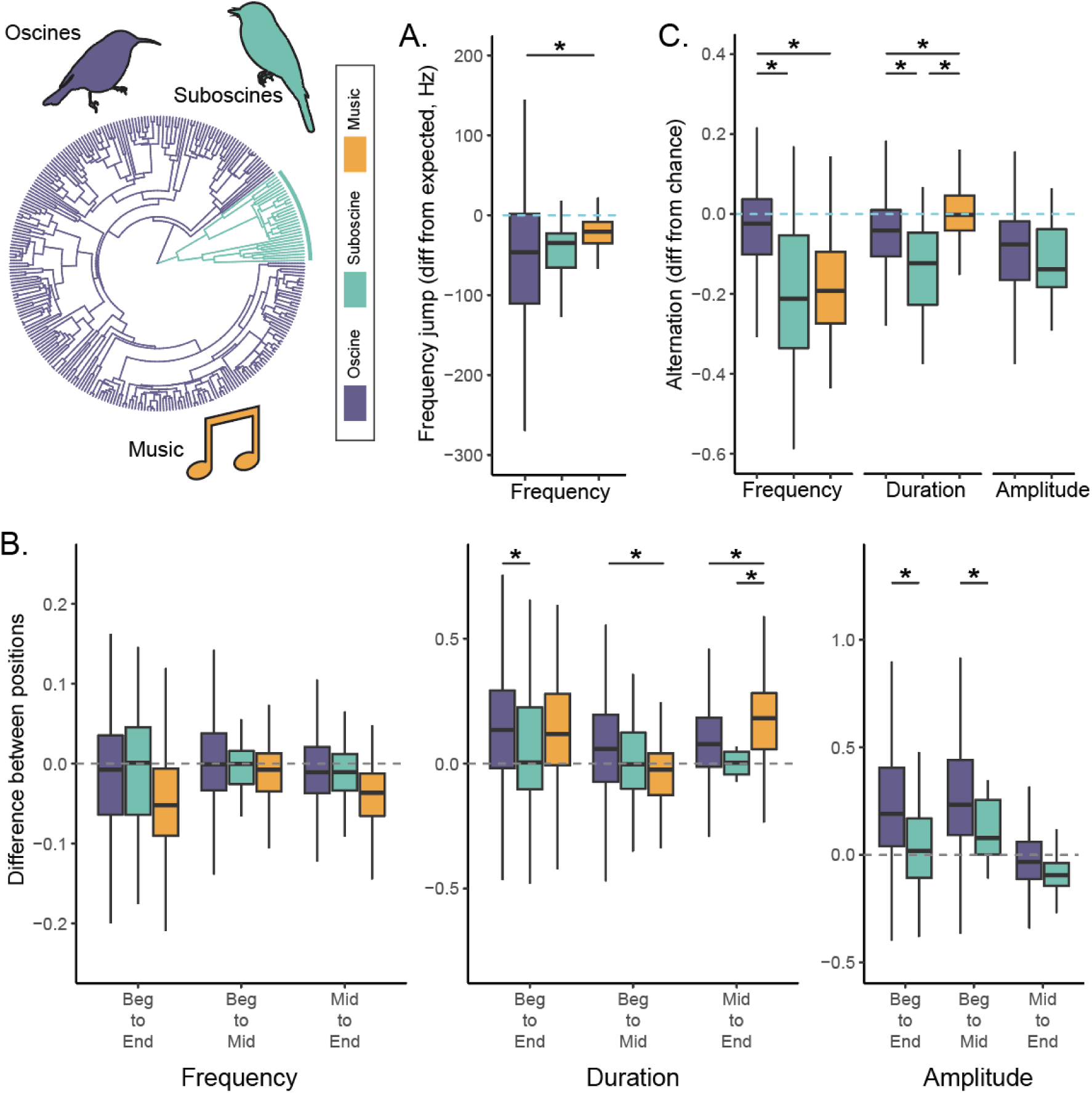
Comparisons of acoustic patterns across oscine, suboscine, and human song. (A) Frequency jumps. Boxplots summarize the differences between observed and expected frequency jumps (negative values indicate smaller jumps than expected), with the dashed line indicating chance. (B) Magnitudes of positional variation in dominant frequency (left), duration (middle), and amplitude (right). Boxplots depict the differences in features between positions, with positive values indicating an increase across positions, negative values indicating a decrease across positions, and the dashed line at zero indicating no change. (C) Degree of alternation in dominant frequency (left), duration (middle), and amplitude (right). Boxplots depict the overall difference in alternation from chance (negative values indicating less alternation than expected), with the dashed line indicating chance. For all panels, oscines are depicted on the left (purple), suboscines in the middle (teal), and music on the right (orange), with all outlier data points removed from the depiction (those beyond 1.5*IQR). Asterisks depict significant differences in patterning between clades (oscines, suboscines, and humans; see Figures 2-4 for analysis of overall patterns).

In general, there were some differences in the magnitude of positional variation in acoustic features between oscines and suboscines (Figure 5B; Table S3). Although there was not significant variation between clades in the direction or degree of position variation for dominant frequency (*p* > 0.8 for all contrasts), significant variation in patterning for syllable duration and amplitude were observed between oscines and suboscines (interaction: *p* < 0.05 for each). Relative to suboscines, oscines demonstrated larger increases in duration from beginning to end syllables and larger changes in syllable amplitude from beginning syllables to both middle and end syllables.

Alternations in all three features were significantly less prevalent than expected by chance for both oscines and suboscines (*p* < 0.001 for all; Figure 5C). However, the degrees of alternation in syllable frequency and duration were significantly different across clades (interaction: *p* < 0.001 for each), driven by the fact that alternation was significantly less prevalent in suboscines than in oscines.

### Comparisons to global patterns in music

While the patterns observed in the vocalizations of birds are broadly similar to patterns described in previous studies of music (Sundberg & Verrillo, 1980; Huron, 1996; Tierney *et al*., 2011), it is important to directly compare the magnitude of these patterns across passerine song and music. To this end, we first extracted and analyzed features from all 118 songs within the Natural History of Song, an open-access corpus that contains songs from around the world (Mehr *et al*., 2019), and then compared the type and magnitude of patterns across oscine, suboscine, and human song. Broadly speaking and consistent with previous analyses of music, we observed that frequency jumps were smaller than expected by chance (*t*_117_ = −11.05, *p* < 0.001) and that alternation in frequency was less prevalent than expected by chance (*t*_114_ = −15.3, *p* < 0.001). We also observed positional variation in frequency (*F*_2,234_ = 42.1, *p* <0.001) and duration (*F*_2,234_ = 58.5, *p* < 0.001), with end notes being longer and lower in frequency than beginning or middle notes (Tukey’s tests: *p* < 0.001 for each) and with beginning notes being longer than middle notes (Tukey’s test: *p* = 0.040). Observed alternation in duration was not different from that expected by chance (*t*_112_ = −0.6, *p* = 0.559).

Finally, we directly compared acoustic patterns in human music to those from oscines and suboscines. Despite the phylogenetic distances and differences in sound production, positional variation in frequency was comparable in magnitude across the three clades (i.e., no significant interaction between group and position: *F*_4,1221_ = 1.8, *p* = 0.121; Figure 5B). Some acoustic patterns were more similar between the two groups of vocal learners: the magnitude of alternation in syllable/note duration (*F*_2,407_ = 20.9, *p* < 0.001) differed across groups, with deviations from chance being comparable between oscines and humans and both significantly different from suboscines. Interestingly, there were acoustic patterns that were more similar between suboscines and humans than between oscines and humans. The magnitude of alternation in frequency differed across groups (*F*_2,409_ = 68.7, *p* < 0.001), with frequency alternation (relative to chance) were comparable between suboscines and humans (music) and both significantly larger (more negative) for than for oscines (*p* < 0.0001 for both).

In addition to these patterns across groups, we observed other differences between human, oscine, and suboscine song. Patterns of positional variation in duration differed across groups (group x position: *F*_4,1221_ = 8.8, *p* < 0.001), with larger increases in duration from beginning to middle elements in oscine song compared to music but larger increases in duration from middle to end elements in music compared to both oscines and suboscines (Tukey’s tests: *p* < 0.02 for all; Figure 5B). Additionally, the degree to which observed frequency jumps deviated from expected differed across groups (*F*_2,412_ = 8.6, *p* < 0.001) with oscines demonstrating a significantly larger (negative) deviation from expected than humans (Tukey’s test: *p* = 0.001; Figure 5A).

## Discussion

Using a large-scale, comparative analysis across hundreds of species, we identified common patterns in vocal sequencing among passerines, focusing on acoustic patterns that are prevalent in voiced music. We found significant patterns in frequency jumps, positional variation, and acoustic alternation within passerine song that resemble those found in music (Huron, 1996; Tierney *et al*., 2011; Savage *et al*., 2015). For example, both oscines (birds that learn their songs) and suboscines (birds that do not learn their songs) sequence the acoustic elements (“syllables”) in their songs in a way that minimizes the change in frequency, a phenomenon that resembles small note intervals across vocal music in humans. In addition, syllables produced at the end of passerine songs tend to be longer and lower in frequency, a pattern that resembles phrase-final elongation and descending pitch contours commonly observed in music.

### Comparisons between passerines and humans

These broad and comparative data enable tests of specific hypotheses regarding the emergence of acoustic universals. For example, because vocalizations in birds and humans require the coordination of respiratory and vocal muscles (Tierney *et al*., 2011; Hutchins, Larrouy-Maestri & Peretz, 2014; Schmidt & Goller, 2016; Goller, 2022a) and share mechanisms of sound production (Elemans *et al*., 2015), it has been proposed that general mechanisms of sound production could lead to acoustic patterns that are widely shared across passerines and humans. Consistent with this hypothesis, positional variation in frequency was similar across oscines, suboscines, and humans, with all three clades showing comparable descending pitch contours. However, many acoustic patterns seemed to differ across oscines, suboscines, and humans, offering relatively limited support for a contribution of broadly shared production mechanisms.

Given that both oscine and human songs are learned and that previous studies have found common patterns between oscine and human song (Baptista & Keister, 2005; Tierney *et al*., 2011), it is possible that vocal learning could contribute to the emergence of acoustic patterns. In this respect, acoustic patterning could be more similar between oscines and humans than between suboscines and humans. Indeed, the magnitudes of alternation and positional variation in syllable durations for oscines were more similar to those in humans than for suboscines, suggesting that learning mechanisms could promote this increased complexity of patterning.

However, contrary to the vocal learning hypothesis, we also found patterns where suboscines were more similar to humans than oscines were. Specifically, the extents of both frequency jumps and frequency alternation were similar between suboscine and human song and were both significantly different from oscine song. These similarities in acoustic patterns highlight potential similarities in vocal production mechanisms between suboscines and humans. Despite that the vocalizations of birds are regulated by bilateral sets of membranes and muscles (syrinx) whereas human vocalizations are regulated by a single set of membranes and muscles (larynx; (Suthers, Goller & Pytte, 1999; Amador *et al*., 2008; Riede & Goller, 2010; Secora *et al*., 2012; Goller, 2022b), some analyses of song production in suboscines have led to the proposal that, in contrast to oscines, suboscines might not be able to independently regulate their bilateral syringeal muscles (Goller *et al*., 2021). In this respect, vocal control in suboscines might be more similar to that in humans. These data emphasize that more extensive inquiries into the patterns and mechanisms of song production in suboscines are required (Goller *et al*., 2021).

Finally, we also observed patterns that were significantly different between passerines and humans. For example, despite that phrase-final elongation was observed in both passerines and humans, human song demonstrated stronger phrase-final elongation than both oscines and suboscines. Phrase-final lengthening in humans has been postulated to be related to changes in muscle control of phonation; in particular, the elongation is thought to be caused by a relaxation of the articulators, causing vocalizations at the end of a phrase to “trail off” (Tierney *et al*., 2011). Interestingly, patterns of respiration differ between avian and human song production; of particular importance is that birds inspire between syllables, taking “mini-breaths” during the silent gaps within song (Schmidt & Goller, 2016; Goller, 2022a). This suggests that the relatively infrequent inspiration when humans sing (or speak) promotes greater elongation of sounds at the end of phrases.

### Comparisons among passerine species

Despite overall similarities in acoustic patterning within passerines, we document substantial variation in acoustic patterns among passerine species. For example, despite the many passerine species that produce songs with phrase-final lengthening, many species within Passerellidae (New World Sparrows) produce songs in which end syllables are shorter in duration than beginning or middle syllables (e.g., white-crowned sparrows produce long whistles at the beginning of their songs; (Rose *et al*., 2004). Compared to other passerines, species within Parulidae (New World Warblers like prairie warblers) demonstrate particularly striking positional variation in amplitude as well as minimal alternation in amplitude (Byers, Akresh & King, 2016). Interspecific diversity is not surprising given the breadth of species of passerines and that vocal learning evolved ∼50 million years ago in this clade (Oliveros *et al*., 2019). Such variation motivates investigations into the evolutionary and ecological factors that shape the nature and strength of these features.

We also discovered differences in alternation, positional variation, and frequency jumps between oscines and suboscines. For example, we observed that suboscines produce songs with significantly less alternation in syllable duration and frequency than oscines. This difference is due, in part, to the greater prevalence of consistently increasing or consistently decreasing acoustic patterns in suboscine song. Relatedly, the magnitude of frequency jumps tended to be smaller in suboscines than oscines. These data are consistent with previous analyses revealing greater frequency ranges and frequency modulation in oscine species than in suboscines (Goller *et al*., 2021) and support the notion that suboscines produce songs with more rigid and simple sequence structure than oscines. Because vocal learning is associated with greater vocal complexity (Nowicki & Searcy, 2014), the decreased level of complexity in acoustic patterning in suboscines may relate to the lack of vocal learning in this group.

Compared to oscines, positional variations in duration and amplitude were weaker in suboscines. Despite the extensive and consistent patterns of phrase-final elongation in oscine song, end syllables are not significantly longer than beginning or middle syllables in suboscine song. This was surprising given shared motor production mechanisms across sub-orders (see also (Tierney *et al*., 2011)). Oscines are also more likely than suboscines to begin their songs with relatively quiet syllables, and it is possible that more oscine species begin their songs with quiet introductory elements than suboscines. The production of introductory elements and changes in respiration before oscine song is thought to reflect motor preparation for learned sequences (Rajan & Doupe, 2013; Méndez, Dukes & Cooper, 2022), and it is possible that more motor preparation is required for optimal performance of oscine song than suboscine song. Given the importance of respiratory regulation for syllable duration and amplitude, these data suggest consistent differences in the control of respiration between oscines and suboscines that may be linked to vocal learning.

### Conclusion

Our expansive and comparative analysis of acoustic patterns in passerine song highlight deep similarities in the organization of passerine and human song. Despite variation in the magnitude of patterns (which could be influenced by variation in phonation mechanisms or vocal learning), passerine and human song widely share descending pitch contours, small frequency jumps, and phrase-final lengthening. In addition to providing extensive evidence of convergent patterns across birds and humans and revealing factors that covary with shared acoustic patterns, our data also highlight surprising similarities between suboscine and human song and underscore the importance of studying vocal mechanisms in clades of birds that do not learn their vocalizations. Given that other movements (i.e., dance) often accompany vocalizations in both passerines and humans (e.g., (Williams, 2001; Dalziell *et al*., 2013; Ota, Gahr & Soma, 2015; Ullrich, Norton & Scharff, 2016; Miles & Fuxjager, 2018)) and that vocal and non-vocal movements are coordinated in humans (reviewed in (Pouw & Fuchs, 2022; Delius & Müller, 2023);), we propose that patterns in non-vocal movements associated with song could also be widely shared across species.

## Methods

### Song acquisition

We aimed to include natural recordings from a wide range of Passerine species, beyond those that have high quality scientific recordings available. As such, we opted to utilize an online, crowd-sourced repository for bird sounds (xeno-canto.org). This resource contains a vast quantity of recordings, with users able to rate the quality of recordings and dispute any mislabeled taxonomic information. These recordings are utilized in a range of scientific works and have confirmed some well-known principles of vocal production [e.g., negative relationship between frequency and body size (Pearse *et al*., 2018)].

To begin, we created a list of species in the order Passeriformes by scraping the “Browse by Taxonomy” section of the website. Then, for each species, we entered a search with the following search terms: (1) the species scientific name (genus and species), (2) “song” as the sound type (as opposed to, e.g., “call”), and (3) the “A” rating (the highest quality, on a scale from A to E, as rated by the recordist). We performed this search using the Xeno-canto API 2.0, which returns a JSON file with details about the recordings that match the search (search date: August 19, 2020). We then used three additional criteria to improve our confidence in the recordings: (1) no background species, (2) the bird was seen, and (3) no playback was used. This information was provided by the recordists when they uploaded the files. We scraped the information from the individual recordings’ webpages. It should be noted that while these checks helped eliminate recordings that would not be appropriate for this project, these checks were not definitive. For example, a recordist might not have listed any background species even though there were background species in the recording, and our manual labeling of files helped to remove such cases from analysis (see below).

We searched randomly on each page of recordings in reverse chronological order and downloaded a maximum of 15 recording files per species for species in which at least five recordings passed our search criteria. Following an initial round of manual annotation on a random subset of the dataset (see below), we focused on species for which we were able to annotate at least one song but had not yet annotated 5 songs.

To analyze sequencing patterns in bird song at the syllable level, we identified song segments, marked the boundaries of each syllable within the songs, and labeled each syllable as being at the beginning, end, or in the middle (i.e., not beginning or end) of the song.

We first identified song segments in the file using an automated procedure adapted from (Oudyk *et al*., 2019). To summarize:

1. We used per-channel energy normalization (PCEN; (Wang *et al*., 2017; Lostanlen *et al*., 2019)) to minimize the background noise in a mel-frequency spectrogram. This method was originally proposed by Google for detection of keywords (e.g., “Hey Google”) in potentially noisy environments, at variable distances from the recording device.
2. We calculated an approximate signal-to-noise ratio (SNR) for each time point in the PCEN spectrogram by subtracting the power of the minimum frequency bin from the power of the maximum frequency bin, dividing by the power of the median frequency bin, then median-smoothing the SNR over 50 ms, giving a SNR curve ranging from 0-1.
3. We performed vocal activity detection with an initial peak threshold on the SNR of (0.25), and then followed the SNR curve in both directions to where it crossed the activity threshold of (0.8). These two crossings were taken as the onset and offset time for each detected sound activity.
4. We stipulated that a gap of 500 ms was needed for two segments to be considered separate songs. Thus, we merged segments that were within 500 ms of each other, and we excluded segments that were within 500 ms of the beginning or the end of the original recording file.
5. In recognition of the limitations of this automated process, we added a 1-second buffer before and after the detected sound activity, so that the human annotators could later judge whether this automated segmentation method captured the beginnings and ends of the songs.

This process, illustrated in Figure S1 resulted in 1,583 audio segments from 299 species that were subsequently deemed labelable by annotators (see below).

#### Passerine song annotation

Songs were semi-automatically segmented into syllables based on an amplitude threshold determined for each file based on recording quality. Thereafter, syllables were manually annotated by individuals with extensive experience with bird song analysis using custom software written in MATLAB (Mathworks, Natick, MA). Song elements (“syllables”) were defined as contiguous epochs of sound (typically > 10 ms, and separated by silence 5 ms of silence; range: 12.3 – 1019.5 ms). The annotators labeled each syllable as being the first syllable, a middle syllable, or the last syllable in a song, with any gap > 500 ms deemed to separate songs. Songs in which the syllables were difficult to extract using the amplitude threshold (e.g., songs with high amplitude background noise or other species vocalizing) were not annotated and, therefore, not included in any analyses. Furthermore, epochs within a song with high background noise or syllables that were difficult to extract were also excluded from analyses (i.e., some songs were only partially analyzed). Individual raters were given random subsets of sound files from across the phylogeny to minimize the influence of any segmentation bias on the overall results. In the end, the dataset consisted of 16.1 ± 10.0 (mean ± SD) songs per species, with some variation across features depending on analysis constraints.

#### Bird song feature extraction

After identifying syllables, we extracted three acoustic features using the Python package Librosa (McFee *et al*., 2015): (1) Duration (length), measured as the number of milliseconds between the onset and offset of the syllable, (2) Dominant frequency (spectral centroid), measured as the frequency (in Hz) with the highest power within each syllable, over frequencies and time, and (3) Amplitude (loudness), measured as the mean root-mean squared value of the waveform (the mean across time). Additionally, we calculated the dominant frequency of the beginning and end of each syllable (first or last 8 ms of syllables). Values for all features were log-transformed for analyses given nonparametric distributions.

#### Analysis of frequency jumps

To assess the change in frequency from one syllable to the next, we calculated the absolute difference in frequency (“jumps”) from the end of one syllable to the beginning of the next syllable (last or first 8 ms) for each consecutive pair of syllables. We then averaged values for each song within a segment, each segment within the same xeno-canto recording, and finally across all recordings from each species (“nested averaging”). This provided an average frequency jump value for each species, and accounts for the fact that multiple songs within the same recording are likely from the same individual.

To assess whether birds sequenced their syllables in a manner that minimized or maximized frequency jumps, we employed Monte Carlo permutations. Specifically, we randomized the order of syllables in each segment 100x and then recalculated the average frequency difference across consecutive syllables. We took the median value from the randomizations, and then conducted the same nested averaging as described above to obtain a mean expected frequency jump for each species. We only included songs with at least 5 consecutive syllables and excluded species without any songs of at least 5 consecutive syllables (*n* = 293 species).

#### Analysis of positional variation in bird song

We measured how our three acoustic features (dominant frequency, duration, and amplitude) varied based on position in the song. For our measurement of position, we compared values for the first syllable in a song (beginning), final syllable in a song (end) and all other syllables (middle). When there were multiple middle syllables in a song, we computed the average value across all middle syllables as the single value for the middle syllable for that segment (Huron, 1996; Tierney *et al*., 2011). We conducted similar nested averaging as described above to obtain an average value for each feature in each position for every species. In some cases, song files ended before the last syllable or started after the first syllable of a song. We only analyzed species for which we had values for all three positions (*n* = 292 species).

#### Analysis of alternation

We next assessed alternation in the three acoustic features. For each set of three consecutive syllables, we assigned a binary value as to whether there was alternation (1) or no alternation (0). For instance, three syllables in a row with durations 10 ms, 20 ms, and 30 ms would be assigned a 0 for no alternation (increase and then increase again), whereas a sequence of 10 ms, 30 ms and 20 ms would be assigned a 1 (increase and then decrease). These binary values were averaged within each song to give an overall level of alternation, and then we conducted the same nested averaging as described above to obtain an average degree of alternation across consecutive syllables for each species. As with our analysis of frequency jumps, we limited our analysis to songs with at least 5 consecutive syllables, and excluded any species that had no songs with at least 5 consecutive syllables (*n* = 293 species).

To compare the observed alternation to that expected by chance, we determined that if any sequence of numbers is randomized, the measurement of alternation we used would produce a value of 0.667 (⅔). For instance, there are 6 possible orders for three syllables with durations of 10 ms, 20 ms and 30 ms (assuming no exact repetition of values). Of these orders, one is up-up, one is down-down, and the other four demonstrate alternation (two up-down and two down-up). This expected value remains the same for songs with more than three different syllable durations (which we confirmed through permutation tests). Thus, observed values above 0.667 would indicate alternation more often than expected by chance.

### Human music

#### Data Acquisition

We used the open dataset of notated songs from around the world, from the ‘Natural History of Song’ project (Mehr *et al*., 2019). Each of the 118 songs were from different parts of the world. This dataset is available at https://doi.org/10.17605/OSF.IO/JMV3Q. We extracted the pitch and duration of each note from the first part (i.e., the top voice) in each song from the transcriptions available in XML format. Based on the way that these pieces were notated, we considered rests and grace notes to mark phrase endings and analyzed each phrase analogous to each bird song as described above. Tied notes were concatenated. The measured pitch of each note (e.g., C5) was converted into frequency (e.g., 523.2 Hz). Durations were measured in beats (a quarter note = 10080). Data on amplitude was not available. Both features were log-transformed for analysis.

We conducted analogous analyses on this dataset to those in bird song. For frequency jumps, we calculated the absolute difference in frequency across adjacent notes and conducted permutation tests to obtain an expected frequency jump given random sequencing. We excluded all cases of repetition when calculating the frequency jump (both in the observed and random datasets). We only calculated the frequency jumps for phrases with at least 3 consecutive notes. We averaged across all phrases within a song to obtain an observed and expected value for each of the 118 songs in the dataset.

For positional variation, our analyses were nearly identical to those in bird song. For each song phrase, we calculated the frequency and duration of the beginning and end notes, as well as the average across all middle notes. We then averaged across all phrases within a song.

Finally, we calculated alternation following a similar protocol as in bird song, but again excluding any cases of note repetition (i.e., notes of the same frequency when calculated frequency alternation, and notes of the same duration when calculating alternation of duration). Like in bird song, the expected level of alternation for any sequence is 2/3 (0.667).

We compared acoustic features extracted from notation for music to those computed from audio recordings of bird song. This is a similar approach to other papers, for example those comparing speech to music (Patel et al., 2006).

### Statistical analyses

We conducted phylogenetically corrected analyses using MCMCglmm models (Hadfield, 2017). Species ID and phylogenetic relatedness were included as random effects. We ran the models for 50,000 iterations, burning the initial 1000 iterations and sampling every 50 iterations thereafter for a total of 998 samples per model. We report the overall Bayesian p-value from the 998 samples (i.e., the percent of samples with the less common direction of effect). All priors were set to uninformative defaults.

We used a phylogenetic tree that was included with the AVONET dataset (Tobias *et al*., 2022), which averaged taxonomic classification from three sources: (BirdLife International, 2020), BirdTree (Jetz *et al*., 2012) and eBird (Clements *et al*., 2021). To visualize features on the phylogenetic trees, we normalized all data to the absolute maximum, setting that value at 1.0 or −1.0, and conducting a subtraction with all other values. There were 22 species with bird song data that were not present in the phylogenetic tree. In 18 cases, we coded the species as its sister or other closely related species. However, in 4 cases, the closest relative(s) was already included in our dataset, thus, we removed these 4 species from analysis.

For frequency jumps, the independent variable was the category of frequency jump (observed or expected), and the dependent variable was with either the average frequency jump observed for the species, or the expected frequency jump computed through randomizations. For positional variation, we included position (beginning, middle, or end) as the independent variable and the mean value of an acoustic feature for each species at each of these positions as the dependent variable. For alternation, the independent variable was the category of alternation (observed or expected), and the dependent variable was either the observed degree of alternation for the species, or the theoretically set expected alternation of 0.667 (see above). We ran similar models to assess differences in patterning between suboscines and oscines by having clade (oscine or suboscine) in full-factorial models.

To assess similar patterns in human music, we conducted linear mixed effects-models (lme4 (Bates *et al*., 2007), with SongID as a random effect. The dependent variables and fixed effects were identical to those described above for the bird song analyses of frequency jumps, positional variation, and alternation. For the positional variation, we conducted Tukey’s tests for post-hoc comparisons among the three positions (beginning, middle, end; package emmeans).

Finally, we ran models to compare patterning in music to that in the oscines and suboscines. For each bird species or each human song, we calculated the (1) difference in frequency jump from expected, (2) differences in frequency and duration across all pairs of positions, and (3) difference in alternation of frequency and duration from expected. For frequency jumps and alternation, we ran simple linear models asking whether group (human, oscine, suboscine) predicted the difference from chance. For positional variation, we ran a full factorial model with comparison (beginning to end, beginning to middle, and middle to end) and group as fixed effects. For all models, we used Tukey’s tests for post-hoc contrasts, but note that we only report effects between music and oscines or suboscines because we used the phylogenetically corrected models described above to calculate differences between the two clades of birds.

## Acknowledgements

We thank A. Vochin and J. Thompson for assistance with data curation and processing. We also thank S.C. Woolley and M.J. Ryan for input on data analysis, presentation, and interpretation. This work was supported by funding from the National Science and Engineering Research Council (#05016 to J.T.S.), Fonds de Recherche du Québec - Nature et technologies (PR-299652 to S.C.W.), Fonds de Recherche du Québec - Santé (K.O.) and the Centre for Research on Brain, Language and Music (J.T.S & L.S.J.). We are deeply indebted to the many individuals that made their data publicly available for use. In particular, to the recordists and maintainers of the xeno-canto project, which continues to amass the largest, to our knowledge, public repository of bird song recordings, and to the Natural History of Song project headed by S. Mehr.

