## Supplementary material for "Pervasive patterns in the songs of passerine birds resemble human music universals and are linked with production and cognitive mechanisms": Table S

### Supplementary information

**Table S1:** List of species in analysis and sample sizes

| Species | Alternation/Jumps | Position | N Sylls | N Songs |
|---|---|---|---|---|
| *Acanthagenys rufogularis* | X | X | 188 | 16 |
| *Acrocephalus aedon* | X | X | 811 | 25 |
| *Acrocephalus agricola* | X | X | 195 | 8 |
| *Acrocephalus australis* | X | X | 110 | 8 |
| *Acrocephalus bistrigiceps* | X | X | 744 | 21 |
| *Acrocephalus dumetorum* | X | X | 185 | 12 |
| *Acrocephalus melanopogon* | X | X | 507 | 16 |
| *Acrocephalus palustris* | X | | 156 | 6 |
| *Acrocephalus schoenobaenus* | X | X | 866 | 25 |
| *Acrocephalus scirpaceus* | X | | 434 | 16 |
| *Acrocephalus stentoreus* | X | X | 215 | 19 |
| *Aegithina tiphia* | X | X | 162 | 17 |
| *Aimophila aestivalis* | X | X | 349 | 23 |
| *Aimophila cassinii* | X | X | 318 | 19 |
| *Aimophila humeralis* | X | X | 134 | 10 |
| *Aimophila rufescens* | X | X | 75 | 13 |
| *Alaemon alaudipes* | X | X | 89 | 11 |
| *Ammodramus bairdii* | X | X | 199 | 8 |
| *Amphispiza bilineata* | X | X | 83 | 5 |
| *Anairetes parulus* | X | X | 420 | 11 |
| *Andropadus importunus* | X | X | 153 | 14 |
| *Anthus godlewskii* | X | X | 190 | 14 |
| *Anthus hodgsoni* | X | X | 226 | 6 |
| *Anthus spinoletta* | X | X | 296 | 9 |
| *Asthenes wyatti* | X | X | 277 | 13 |
| *Atlapetes latinuchus* | X | X | 136 | 25 |
| *Atlapetes pallidiceps* | X | X | 175 | 19 |
| *Attila rufus* | X | X | 397 | 66 |
| *Baeolophus bicolor* | X | X | 37 | 11 |
| *Baeolophus inornatus* | X | X | 171 | 10 |
| *Basileuterus flaveolus* | X | X | 108 | 9 |
| *Basileuterus rufifrons* | X | X | 329 | 18 |
| *Cacicus leucoramphus* | X | X | 32 | 8 |
| *Calandrella brachydactyla* | X | X | 307 | 20 |
| *Calandrella cheleensis* | X | X | 365 | 15 |
| *Calandrella rufescens* | X | X | 146 | 11 |
| *Calcarius pictus* | X | X | 197 | 18 |
| *Camptostoma imberbe* | X | X | 77 | 16 |
| *Campylorhamphus falcularius* | X | X | 139 | 15 |
| *Campylorhynchus brunneicapillus* | X | X | 399 | 18 |
| *Cardinalis sinuatus* | X | X | 24 | 4 |
| *Carduelis cannabina* | X | X | 233 | 13 |
| *Carduelis carduelis* | X | X | 345 | 12 |
| *Carduelis chloris* | X | X | 230 | 20 |
| *Carduelis psaltria* | X | X | 292 | 16 |
| *Carduelis spinus* | X | X | 263 | 16 |

| | | | | |
|---|---|---|---|---|
| *Carduelis tristis* | X | X | 216 | 15 |
| *Carpodacus erythrinus* | X | X | 129 | 24 |
| *Carpodacus mexicanus* | X | X | 344 | 19 |
| *Catharus ustulatus* | X | X | 96 | 15 |
| *Catherpes mexicanus* | X | X | 349 | 23 |
| *Cercomacra melanaria* | X | X | 105 | 24 |
| *Certhia americana* | X | X | 91 | 19 |
| *Certhia brachydactyla* | X | X | 160 | 21 |
| *Certhia familiaris* | X | X | 272 | 16 |
| *Cettia diphone* | | X | 41 | 7 |
| *Cettia fortipes* | | X | 38 | 10 |
| *Cettia vulcania* | | X | 15 | 10 |
| *Cichladusa guttata* | X | X | 175 | 19 |
| *Cinnycerthia olivascens* | X | | 94 | 6 |
| *Cisticola chiniana* | X | X | 169 | 20 |
| *Cistothorus palustris* | X | X | 81 | 5 |
| *Cyanocorax caeruleus* | X | X | 62 | 11 |
| *Cyclarhis gujanensis* | X | X | 106 | 17 |
| *Cyphorhinus thoracicus* | X | | 35 | 7 |
| *Dendroica aestiva* | X | X | 144 | 16 |
| *Dendroica castanea* | X | X | 43 | 6 |
| *Dendroica coronata* | X | X | 240 | 19 |
| *Dendroica discolor* | X | X | 337 | 19 |
| *Dendroica graciae* | X | X | 185 | 13 |
| *Dendroica kirtlandii* | X | X | 77 | 9 |
| *Dendroica striata* | X | X | 112 | 11 |
| *Dendroica tigrina* | X | X | 100 | 20 |
| *Dicaeum trigonostigma* | X | X | 131 | 19 |
| *Dolichonyx oryzivorus* | X | X | 287 | 9 |
| *Drymophila caudata* | X | X | 270 | 41 |
| *Drymophila ochropyga* | X | X | 69 | 28 |
| *Drymophila squamata* | X | X | 305 | 51 |
| *Dumetella carolinensis* | X | X | 161 | 18 |
| *Dysithamnus mentalis* | X | X | 265 | 18 |
| *Emberiza bruniceps* | X | X | 193 | 11 |
| *Emberiza buchanani* | X | X | 102 | 10 |
| *Emberiza cia* | X | X | 283 | 17 |
| *Emberiza cioides* | X | X | 222 | 19 |
| *Emberiza cirlus* | X | X | 404 | 19 |
| *Emberiza citrinella* | X | X | 267 | 22 |
| *Emberiza hortulana* | X | X | 128 | 18 |
| *Emberiza leucocephalos* | X | X | 299 | 24 |
| *Emberiza pusilla* | X | X | 310 | 21 |
| *Emberiza schoeniclus* | X | X | 205 | 29 |
| *Embernagra platensis* | | X | 29 | 8 |
| *Ergaticus ruber* | X | X | 132 | 10 |
| *Erithacus rubecula* | X | X | 248 | 16 |
| *Erythropygia galactotes* | X | X | 85 | 15 |
| *Erythropygia leucophrys* | X | X | 58 | 9 |
| *Ficedula albicollis* | X | X | 217 | 21 |

| | | | | |
|---|---|---|---|---|
| *Ficedula hypoleuca* | X | X | 95 | 9 |
| *Ficedula mugimaki* | X | X | 127 | 7 |
| *Ficedula zanthopygia* | X | X | 38 | 10 |
| *Galerida cristata* | X | X | 218 | 21 |
| *Galerida theklae* | X | X | 554 | 28 |
| *Geothlypis trichas* | X | X | 161 | 15 |
| *Grallaria quitensis* | | X | 93 | 16 |
| *Habia rubica* | X | X | 79 | 13 |
| *Herpsilochmus rufimarginatus* | X | X | 886 | 34 |
| *Heterophasia pulchella* | X | X | 65 | 8 |
| *Hippolais icterina* | X | X | 112 | 7 |
| *Hippolais opaca* | X | X | 454 | 16 |
| *Hippolais pallida* | X | X | 428 | 16 |
| *Hippolais polyglotta* | X | X | 1269 | 23 |
| *Hirundo rustica* | X | X | 124 | 12 |
| *Hylophilus flavipes* | X | X | 172 | 11 |
| *Hypocnemis cantator* | X | X | 44 | 4 |
| *Hypsipetes borbonicus* | X | X | 28 | 5 |
| *Icterus spurius* | X | X | 113 | 6 |
| *Irania gutturalis* | X | X | 136 | 9 |
| *Junco hyemalis* | X | X | 401 | 23 |
| *Lanius excubitor* | X | X | 32 | 5 |
| *Lanius ludovicianus* | X | X | 115 | 12 |
| *Lanius somalicus* | X | X | 70 | 16 |
| *Lichmera indistincta* | X | X | 93 | 11 |
| *Locustella certhiola* | X | X | 550 | 18 |
| *Loxia curvirostra* | X | X | 117 | 11 |
| *Loxia leucoptera* | X | X | 681 | 12 |
| *Lullula arborea* | X | X | 487 | 27 |
| *Luscinia calliope* | X | X | 189 | 14 |
| *Luscinia cyane* | X | X | 120 | 17 |
| *Luscinia megarhynchos* | X | X | 279 | 19 |
| *Luscinia svecica* | X | X | 303 | 16 |
| *Mackenziaena severa* | X | X | 142 | 24 |
| *Malacocincla abbotti* | X | X | 56 | 17 |
| *Mecocerculus leucophrys* | X | X | 269 | 11 |
| *Melanocorypha leucoptera* | X | | 252 | 12 |
| *Melanoptila glabrirostris* | X | X | 51 | 7 |
| *Melanotis caerulescens* | X | X | 91 | 12 |
| *Melospiza georgiana* | X | X | 245 | 8 |
| *Melospiza lincolnii* | X | X | 175 | 11 |
| *Melospiza melodia* | X | X | 358 | 19 |
| *Microrhopias quixensis* | X | X | 240 | 39 |
| *Mimus gilvus* | X | X | 94 | 15 |
| *Mimus patagonicus* | X | X | 130 | 11 |
| *Mimus polyglottos* | X | X | 231 | 21 |
| *Minla strigula* | X | X | 22 | 4 |
| *Mirafra hypermetra* | X | X | 131 | 14 |
| *Monticola saxatilis* | X | X | 222 | 15 |
| *Montifringilla nivalis* | X | X | 94 | 8 |

| | | | | |
|---|---|---|---|---|
| *Muscicapa striata* | | X | 19 | 4 |
| *Myioborus miniatus* | X | X | 135 | 15 |
| *Myrmeciza ferruginea* | X | X | 348 | 32 |
| *Myrmeciza immaculata* | X | X | 93 | 19 |
| *Myrmotherula gularis* | X | X | 86 | 12 |
| *Nectarinia chalybea* | X | X | 163 | 6 |
| *Nectarinia senegalensis* | X | X | 102 | 13 |
| *Odontorchilus cinereus* | X | X | 238 | 10 |
| *Oenanthe hispanica* | X | X | 177 | 16 |
| *Oenanthe leucura* | X | X | 83 | 5 |
| *Oenanthe oenanthe* | X | X | 323 | 25 |
| *Oenanthe pleschanka* | X | X | 199 | 21 |
| *Oporornis agilis* | X | X | 142 | 13 |
| *Oporornis philadelphia* | X | X | 47 | 9 |
| *Oporornis tolmiei* | X | X | 80 | 11 |
| *Oreoscoptes montanus* | X | X | 604 | 15 |
| *Orthotomus sericeus* | X | X | 117 | 14 |
| *Orthotomus sutorius* | X | X | 155 | 16 |
| *Pachycephala nudigula* | X | X | 119 | 12 |
| *Pachycephala pectoralis* | X | X | 104 | 14 |
| *Paradisaea rubra* | X | X | 84 | 5 |
| *Parus ater* | X | X | 210 | 20 |
| *Parus atricapillus* | X | X | 93 | 17 |
| *Parus caeruleus* | X | X | 57 | 5 |
| *Parus cristatus* | X | X | 268 | 15 |
| *Parus cyanus* | X | X | 370 | 25 |
| *Parus gambeli* | X | X | 203 | 24 |
| *Parus major* | X | X | 489 | 42 |
| *Parus montanus* | X | X | 174 | 26 |
| *Parus monticolus* | X | X | 98 | 14 |
| *Parus nuchalis* | X | X | 93 | 13 |
| *Parus palustris* | X | X | 237 | 23 |
| *Passer hispaniolensis* | X | X | 57 | 7 |
| *Passerella iliaca* | X | X | 861 | 72 |
| *Passerina amoena* | X | X | 337 | 25 |
| *Passerina caerulea* | X | X | 218 | 15 |
| *Passerina cyanea* | X | X | 102 | 9 |
| *Percnostola rufifrons* | X | X | 253 | 32 |
| *Perisoreus infaustus* | X | X | 36 | 5 |
| *Phaeomyias murina* | X | X | 148 | 16 |
| *Phoenicurus ochruros* | X | X | 73 | 9 |
| *Phoenicurus phoenicurus* | X | X | 256 | 22 |
| *Phrygilus fruticeti* | X | X | 23 | 5 |
| *Phylloscartes ventralis* | X | X | 187 | 8 |
| *Phylloscopus affinis* | X | X | 90 | 14 |
| *Phylloscopus bonelli* | X | X | 240 | 21 |
| *Phylloscopus borealis* | X | X | 287 | 13 |
| *Phylloscopus claudiae* | X | X | 162 | 11 |
| *Phylloscopus coronatus* | X | X | 86 | 17 |
| *Phylloscopus davisoni* | X | X | 124 | 15 |

| | | | | |
|---|---|---|---|---|
| *Phylloscopus fuscatus* | X | X | 50 | 7 |
| *Phylloscopus inornatus* | X | X | 43 | 10 |
| *Phylloscopus proregulus* | X | X | 489 | 25 |
| *Phylloscopus reguloides* | X | X | 195 | 19 |
| *Phylloscopus schwarzi* | X | X | 202 | 14 |
| *Phylloscopus trochilus* | X | X | 351 | 21 |
| *Pipilo chlorurus* | X | X | 269 | 20 |
| *Pipilo maculatus* | X | X | 135 | 11 |
| *Pipraeidea melanonota* | X | X | 62 | 8 |
| *Piranga leucoptera* | X | X | 21 | 4 |
| *Piranga olivacea* | X | X | 63 | 11 |
| *Pitangus sulphuratus* | X | X | 347 | 86 |
| *Plectrophenax nivalis* | X | X | 265 | 22 |
| *Ploceus baglafecht* | X | X | 22 | 3 |
| *Ploceus cucullatus* | X | X | 43 | 5 |
| *Podoces pleskei* | X | X | 396 | 14 |
| *Polioptila dumicola* | X | X | 118 | 16 |
| *Pooecetes gramineus* | X | X | 110 | 7 |
| *Prinia flaviventris* | X | X | 141 | 13 |
| *Prinia gracilis* | X | X | 147 | 17 |
| *Prinia subflava* | X | X | 110 | 10 |
| *Prosthemadera novaeseelandiae* | X | | 23 | 3 |
| *Prunella fulvescens* | X | X | 281 | 16 |
| *Prunella modularis* | X | X | 404 | 23 |
| *Pseudoleistes guirahuro* | X | X | 30 | 4 |
| *Pteruthius flaviscapis* | X | X | 77 | 14 |
| *Pygochelidon cyanoleuca* | X | X | 27 | 5 |
| *Pyriglena leuconota* | X | X | 805 | 44 |
| *Pyriglena leucoptera* | X | X | 169 | 22 |
| *Regulus calendula* | X | X | 381 | 18 |
| *Regulus ignicapilla* | X | X | 287 | 19 |
| *Regulus regulus* | X | X | 242 | 16 |
| *Regulus satrapa* | X | X | 159 | 15 |
| *Remiz coronatus* | X | X | 30 | 9 |
| *Rhipidura leucophrys* | X | X | 76 | 8 |
| *Salpinctes obsoletus* | X | X | 180 | 14 |
| *Saltator atricollis* | X | X | 201 | 21 |
| *Saltator aurantiirostris* | X | X | 55 | 10 |
| *Saltator coerulescens* | X | X | 55 | 10 |
| *Saltator grossus* | X | X | 43 | 12 |
| *Saltator similis* | X | X | 106 | 17 |
| *Saltator striatipectus* | X | X | 57 | 9 |
| *Saxicola rubetra* | X | X | 55 | 7 |
| *Schiffornis virescens* | X | X | 238 | 53 |
| *Scotocerca inquieta* | X | X | 376 | 17 |
| *Seiurus noveboracensis* | X | X | 129 | 11 |
| *Sericossypha albocristata* | X | X | 44 | 9 |
| *Serinus mozambicus* | X | X | 81 | 15 |
| *Sitta canadensis* | X | X | 399 | 11 |
| *Sitta pusilla* | X | X | 115 | 15 |

| | | | | |
|---|---|---|---|---|
| *Spindalis zena* | X | X | 445 | 14 |
| *Spiza americana* | X | X | 30 | 5 |
| *Spizella arborea* | X | X | 169 | 13 |
| *Spizella passerina* | X | X | 806 | 22 |
| *Spizella pusilla* | X | X | 111 | 5 |
| *Spizella wortheni* | X | X | 249 | 14 |
| *Sporophila caerulescens* | X | X | 139 | 15 |
| *Sporophila torqueola* | X | X | 386 | 22 |
| *Sturnella bellicosa* | X | X | 79 | 13 |
| *Sturnella loyca* | X | X | 52 | 7 |
| *Sturnella magna* | X | X | 109 | 20 |
| *Sturnella neglecta* | X | X | 151 | 17 |
| *Sylvia atricapilla* | X | X | 375 | 16 |
| *Sylvia cantillans* | X | X | 907 | 35 |
| *Sylvia conspicillata* | X | X | 282 | 16 |
| *Sylvia curruca* | X | X | 458 | 18 |
| *Sylvia hortensis* | X | X | 500 | 39 |
| *Sylvia nisoria* | X | X | 508 | 19 |
| *Sylvia undata* | X | X | 398 | 22 |
| *Thamnophilus caerulescens* | X | X | 126 | 16 |
| *Thamnophilus doliatus* | X | X | 844 | 36 |
| *Thryomanes bewickii* | X | X | 292 | 18 |
| *Thryothorus ludovicianus* | X | X | 241 | 17 |
| *Thryothorus sclateri* | X | X | 38 | 7 |
| *Tiaris olivaceus* | X | X | 223 | 12 |
| *Toxostoma bendirei* | X | X | 389 | 12 |
| *Toxostoma crissale* | X | X | 143 | 15 |
| *Toxostoma curvirostre* | X | X | 181 | 15 |
| *Toxostoma lecontei* | X | | 76 | 7 |
| *Toxostoma redivivum* | X | X | 207 | 17 |
| *Troglodytes troglodytes* | X | X | 1444 | 30 |
| *Turdus amaurochalinus* | X | X | 121 | 20 |
| *Turdus flavipes* | X | X | 107 | 13 |
| *Turdus grayi* | X | X | 91 | 11 |
| *Turdus iliacus* | X | X | 114 | 11 |
| *Turdus leucomelas* | X | X | 102 | 13 |
| *Turdus leucops* | X | X | 76 | 9 |
| *Turdus merula* | X | X | 102 | 12 |
| *Turdus philomelos* | X | X | 254 | 28 |
| *Turdus rufiventris* | X | X | 570 | 30 |
| *Turdus torquatus* | X | X | 21 | 6 |
| *Turdus viscivorus* | X | X | 37 | 6 |
| *Tyrannus melancholicus* | X | X | 589 | 47 |
| *Vermivora luciae* | X | X | 286 | 20 |
| *Vermivora peregrina* | X | X | 348 | 12 |
| *Vermivora ruficapilla* | X | X | 149 | 11 |
| *Vireo atricapilla* | X | X | 99 | 10 |
| *Vireo bellii* | X | X | 267 | 25 |
| *Vireo gilvus* | X | X | 215 | 19 |
| *Wilsonia pusilla* | X | X | 228 | 23 |

| | | | | |
|---|---|---|---|---|
| *Xenodacnis parina* | X | X | 27 | 5 |
| *Xiphorhynchus fuscus* | X | X | 72 | 5 |
| *Zonotrichia leucophrys* | X | X | 109 | 14 |

**Table S2:** Model outputs

Sequencing patterns in birdsong. Reported is the mean and 95% confidence interval for each coefficient across all samples from MCMCglmm analyses as well as the Bayesian p-value (see Methods). Pagel's λ and Blomberg's K are reported along with their associated p-values obtained from likelihood or randomization tests, respectively, based on average data per species.

| *Analysis* | *Feature* | *Contrast* | *Coefficient [95% CI]* | *p* | *λ* | *p (λ)* | *K* | *p (K)* |
|---|---|---|---|---|---|---|---|---|
| *Jumps* | Frequency | chance | –66.78 [–80.33, –52.00] | < 0.001 | 0.12 | 0.002 | 0.002 | 0.746 |
| *Position* | Frequency | beg to end | –0.0072 [–0.0154, –0.0003] | 0.071 | 0.04 | 0.743 | 0.028 | 0.248 |
| | | beg to mid | 0.0026 [–0.0056, 0.0105] | 0.553 | < 0.001 | 1 | 0.021 | 0.29 |
| | | mid to end | –0.0098 [–0.0169, –0.0027] | 0.006 | 0.146 | 0.336 | 0.13 | 0.01 |
| | Duration | beg to end | 0.1378 [0.1119, 0.1654] | < 0.001 | 0.293 | 0.031 | 0.101 | 0.047 |
| | | beg to mid | 0.0496 [0.0219, 0.0720] | < 0.001 | 0.339 | 0.007 | 0.153 | 0.001 |
| | | mid to end | 0.0882 [0.0654, 0.1103] | < 0.001 | < 0.001 | 1 | 0.087 | 0.066 |
| | Amplitude | beg to end | 0.2161 [0.1874, 0.2437] | < 0.001 | 0.704 | < 0.001 | 0.005 | 0.534 |
| | | beg to mid | 0.2487 [0.2202, 0.2764] | < 0.001 | 0.631 | < 0.001 | 0.002 | 0.813 |
| | | mid to end | –0.0326 [–0.0563, –0.0078] | 0.028 | 0.603 | < 0.001 | 0.004 | 0.587 |
| *Alternation* | Frequency | chance | –0.0517 [–0.0654, –0.0379] | < 0.001 | 0.23 | < 0.001 | < 0.001 | 0.963 |
| | Duration | chance | –0.0644 [–0.0773, –0.0526] | < 0.001 | 0.22 | < 0.001 | 0.005 | 0.544 |
| | Amplitude | chance | –0.0970 [–0.1080, –0.0841] | < 0.001 | 0.23 | 0.025 | 0.001 | 0.898 |

**Table S3:** Model outputs for oscine/suboscine analyses
Variation in acoustic patterning based on clade (oscine vs suboscine). Reported is the mean and 95% confidence interval for each coefficient (from MCMCglmm) and Bayesian p-values.

| *Analysis* | *Feature* | *Contrast* | *Coefficient [95% CI]* | *p* |
|---|---|---|---|---|
| *Jumps* | Frequency | expected | –69.02 [–84.30, –54.54] | < 0.001 |
| | | Clade | –246.7 [–495.3, 5.4] | 0.069 |
| | | Clade * expected | 24.45 [–20.16, 71.65] | 0.333 |
| *Position* | Frequency | beg to end | –0.0075 [–0.0165, 0.0012] | 0.098 |
| | | beg to mid | 0.0022 [–0.0072, 0.0100] | 0.596 |
| | | mid to end | –0.0099 [–0.0172, –0.0022] | 0.033 |
| | | Clade | –0.0267 [–0.1866, 0.1147] | 0.724 |
| | | Clade * beg to end | 0.0030 [–0.0250, 0.0319] | 0.841 |
| | | Clade * beg to mid | 0.0022 [–0.0263, 0.0282] | 0.896 |
| | | Clade * mid to end | –0.0008 [–0.0245, 0.0241] | 0.945 |
| | Duration | beg to end | 0.1463 [0.1199, 0.1767] | < 0.001 |
| | | beg to mid | 0.0512 [0.0231, 0.0770] | < 0.001 |
| | | mid to end | 0.0945 [0.0713, 0.1192] | < 0.001 |
| | | Clade | 0.1287 [–0.0989, 0.3657] | 0.276 |
| | | Clade * beg to end | –0.0944 [–0.1909, –0.0185] | 0.037 |
| | | Clade * beg to mid | –0.0313 [–0.1231, 0.0580] | 0.508 |
| | | Clade * mid to end | 0.0631 [–0.0147, 0.1424] | 0.188 |
| | Amplitude | beg to end | 0.2347 [0.2027, 0.2618] | < 0.001 |
| | | beg to mid | 0.2628 [0.2346, 0.2933] | < 0.001 |
| | | mid to end | –0.0281 [–0.0533, –0.0026] | 0.069 |
| | | Clade | 0.1803 [–0.0290, –0.3669] | 0.09 |
| | | Clade * beg to end | –0.1878 [–0.2759, –0.0994] | < 0.001 |
| | | Clade * beg to mid | –0.1440 [–0.2236, –0.0450] | < 0.001 |
| | | Clade * mid to end | 0.04370 [–0.0297, 0.1162] | 0.329 |
| *Alternation* | Frequency | expected | –0.0359 [–0.0493, –0.0197] | < 0.001 |
| | | Clade | 0.0283 [–0.0437, 0.0983] | 0.435 |
| | | Clade * expected | –0.1720 [–0.2224, –0.1327] | < 0.001 |
| | Duration | expected | –0.0563 [–0.0690, –0.0446] | < 0.001 |
| | | Clade | 0.0151 [–0.0460, 0.0876] | 0.653 |
| | | Clade * expected | –0.0886 [–0.1256, –0.0447] | < 0.001 |
| | Amplitude | expected | –0.0964 [–0.1083, –0.0835] | < 0.001 |
| | | Clade | 0.0186 [–0.0461, 0.0850] | 0.567 |
| | | Clade * expected | –0.0010 [–0.0504, 0.0288] | 0.624 |

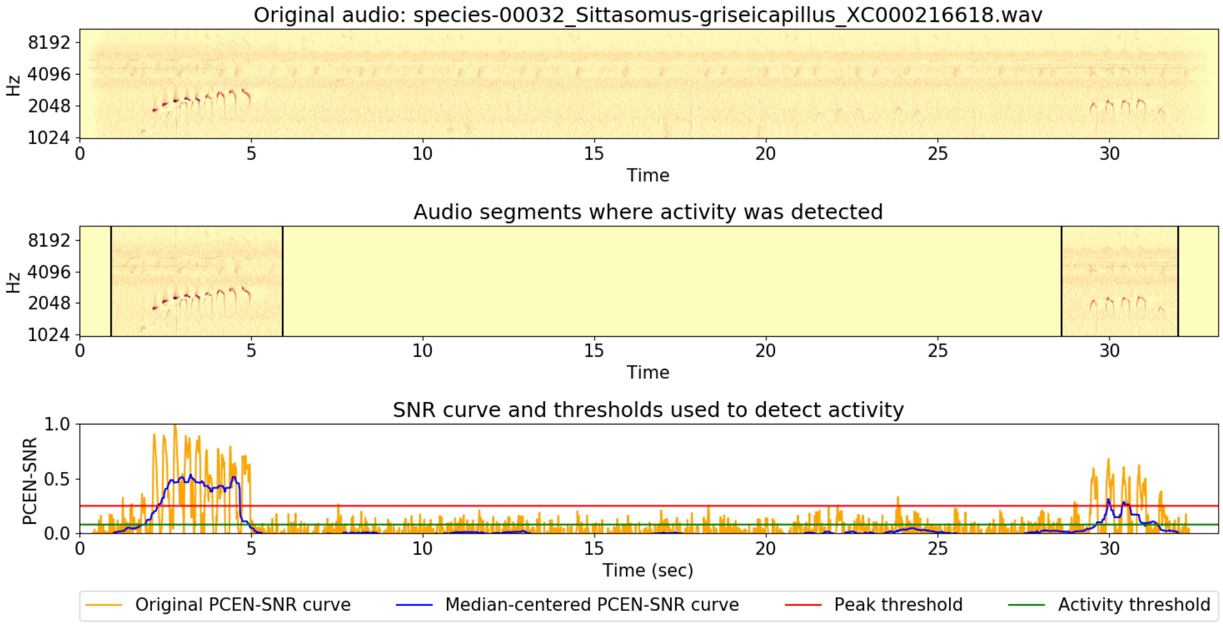


***Figure S1.*** *Illustration of the process of automatically selecting song segments (middle panel) from full recordings (top panel), using the signal-to-noise ratio (SNR) curves and the peak and activity thresholds (bottom panel; Oudyk et al., 2019). Annotators were presented with segments where activity (sound) was detected (i.e., the 'songs' in the middle panel). This is an example for* Sittasomus griseicapillus, *a suboscine.*
